# Ochratoxin A induces behavioral and neurochemical changes in adult zebrafish

**DOI:** 10.1101/2021.10.18.464868

**Authors:** Jéssica Valadas, Adrieli Sachett, Matheus Marcon, Leonardo M. Bastos, Angelo Piato

**Author notes:** Correspondence to: Angelo Piato, Ph.D. Programa de Pós-graduação em Ciências Biológicas: Farmacologia e Terapêutica, Instituto de Ciências Básicas da Saúde, Universidade Federal do Rio Grande do Sul (UFRGS), Avenida Sarmento Leite, 500/305, Porto Alegre, RS 90050-170, Brazil.

## Abstract

Ochratoxin A (OTA) is a mycotoxin produced by species of filamentous fungi widely found as a contaminant in food and with high toxic potential. Studies have shown that this toxin cause kidney and liver damage, however, data on the effects of exposure to OTA on the central nervous system are still scarce. Zebrafish (*Danio rerio*) is a teleost often used in translational research due to its physiological, genetic, and behavioral homology with mammals, in addition to being useful as an environmental bioindicator. Thus, this study aimed to investigate the effects of exposure to OTA on behavioral and neurochemical parameters in adult zebrafish. The animals were treated with different doses of OTA (1.38, 2.77, and 5.53 mg/kg) and submitted to behavioral evaluations in the open tank and social interaction tests. Subsequently, they were euthanized, and the brains were used to assess markers associated with oxidative status. In the open tank test OTA induced changes in distance, absolute turn angle, mean speed, and time-freezing. However, no significant effects were observed in the social interaction test. Moreover, OTA also induced alterations in neurochemical parameters with changes in non-protein thiols (NPSH), glutathione peroxidase (GPx), glutathione-S-transferase (GST), and glutathione reductase (GR). This study showed that OTA can affect neurobiological aspects in zebrafish even at low doses.

## INTRODUCTION

It is estimated that 200 thousand people are added daily to the world’s demand for food (Nellemann et al., 2009). With the projections that by 2050 the world will reach 9.8 billion inhabitants (United Nations, 2017), the search for solutions to meet these needs becomes urgent. Currently, the tools used to solve this issue are responsible for creating other problems. For example, the increase of pesticides in large crops is already causing serious environmental and public health impacts (Langley and Mort, 2012; Rani et al., 2021; World Health Organization, 2006); Improperly tampering with livestock products has become a crime that puts consumers’ lives in danger (Cavin et al., 2018; Xin and Stone, 2008); Industry investment in processed foods has been linked to the incidence of obesity, diabetes, celiac disease and heart disease (Aguayo-Patrón and Calderón de la Barca, 2017; Anand et al., 2015; Canella et al., 2014). Although for a long time, environmental conditions and inadequate storage of food products have been ignored, today it is already clear that these conducts are responsible for the increasing presence of mycotoxins (Marroquín-Cardona et al., 2014).

Mycotoxins are naturally occurring compounds in species of fungi and are potentially toxic (Tola and Kebede, 2016). Ochratoxin A (OTA) is a mycotoxin produced by filamentous fungi and belongs to the ochratoxin subgroup, along with ochratoxin B and C. However, OTA has more natural occurrence and higher toxicity than other ochratoxins. OTA has become a very common contaminant in food and the ecosystem. There is evidence of the presence of OTA in water sources (Hu et al., 2017; Mata et al., 2015) and sea animals (Sun et al., 2015). The highest incidence of detection, however, occurs in food. OTA has already been found in many types of food in the world, including meats found in Croatia (Pleadin et al., 2015), in Brazilian and European coffee (Almeida et al., 2007; v. d. Stegen et al., 1997), wines, and beers from Chile and Hungary (VARGA et al., 2014; Vega et al., 2012), fruits in Argentina and Canada (Lombaert et al., 2004; Magnoli et al., 2004), European juices (Jørgensen, 2005) and several other types of products across the globe.

The exportation market moves around billions of dollars per year (Food and Agriculture Organization (FAO), 2019b) and billions of food tons (Food and Agriculture Organization (FAO), 2019a) are transferred from some places in the world and transported to other countries with different laws and cultures. Most of those nations have protocols and specific regulations for the limits tolerable of contaminants in the food, including OTA (Bureau of Chemical Safety et al., 2009; Ministério da Saúde and Agência Nacional de Vigilância Sanitária, 2011; Official Journal of the European Union, 2006). However, there is no consensus on acceptable limits for this contaminant among countries and, as the trade develops, the OTA present in food crosses borders and easily spreads around the world due to lack of consent between health inspection standards.

The mechanism throughout OTA toxicity it’s not clear yet. It is believed to be related to inhibition of proteins synthesis caused by the competition between the phenylalanine group of OTA and phenylalanine amino acid. Other pathways such as cellular energy production and genetic are also affected (Kőszegi and Poór, 2016). The effects of OTA have already been evaluated in rodents (Castegnaro et al., 1998; Kanisawa and Suzuki, 1978), birds (Stoev, 2010), and fish (Doster et al., 1974; Manning et al., 2003). The toxin has been associated with immune modulation (Lea et al., 1989), hepatic (Qi et al., 2015), and kidney diseases (Abid et al., 2003; Fuchs and Peraica, 2005). OTA has also been increasingly associated with neuropsychiatric diseases (Brewer et al., 2013; V Sava et al., 2006a, 2006c; Yoon et al., 2009), however, despite the importance of these reports, there is still little information regarding the behavioral and neurochemical effects related to OTA on non-target organisms. Therefore, OTA is an important contaminant for both environment and food commodities, but there are still several gaps in the knowledge about the effects of that toxin in organisms.

Native from Asia, the zebrafish is a teleost that has high genetic and physiological homology with humans (Lieschke and Currie, 2007), by this reason, this species has been used as a research animal model for different lines such as embryology and development (Hao et al., 2013; Keller et al., 2008), oxidative stress (Choi et al., 2010; Marcon et al., 2018), behavior (Abozaid et al., 2020; Nabinger et al., 2021; Reis et al., 2020) and genetic (Nasevicius and Ekker, 2000; Falcão et al., 2021). Moreover, this aquatic animal is a very interesting environmental bioindicator used in toxicology and ecotoxicology research, due to its capacity to simulate the conditions of an animal in its natural ecosystem (Asharani et al., 2008; Park et al., 2020; Valadas et al., 2019). In this context, since zebrafish is a suitable environmental bioindicator used in toxicology research, this study aimed to investigate the behavior and neurochemical effects of OTA in adult zebrafish.

## MATERIALS AND METHODS

### Animals

The experiments were performed using 96 adult short-fin wild-type zebrafish (*Danio rerio*, Hamilton, 1822) of both sexes (50:50 male:female ratio) obtained from the local commercial supplier were used at the protocols. The animals were housed in a maximum density of two fish per liter of water in 16-L tanks (40 × 20 × 24 cm) and under a 14–10-h day/night cycle for 10 days before any procedure. Water parameters such as pH (7.0 ± 0.3), chlorine, ammonia (< 0.01 mg/L), and temperature (26 °C ± 2) were controlled. Fish were fed twice a day with commercial flake food (Poytara^®^, Brazil) and supplementation of brine shrimp (*Artemia salina*). After the behavior tests, the animals were euthanized by hypothermic shock (2 °- 4 °C) followed by decapitation, according to the AVMA Guidelines for the Euthanasia of Animals (Leary and Johnson, 2020). All procedures were approved by the Universidade Federal do Rio Grande do Sul ethical committee (#37761/2020). The protocols were reported following ARRIVE Guidelines 2.0 (Percie du Sert et al., 2020).

### Drugs

Ochratoxin A (OTA) (CAS 303-47-9), dimethyl sulfoxide (DMSO) (CAS 67-68-5), and tricaine (MS-222) (CAS 886–86-2) were obtained from Sigma-Aldrich (St. Louis, MO, USA). Sodium chloride solution 0.9% (saline, ADV Farma, SP, Brazil) was obtained from a local commercial supplier. OTA was dissolved into DMSO (final concentration of 10% DMSO). The OTA doses were based on the LD50 for intraperitoneal injection on rainbow trout (*Salmo gairdneri* or *Oncorhynchus mykiss*) (Doster et al., 1972) since there are no similar studies on adult zebrafish. The established doses for OTA in this study were 1.38, 2.77, and 5.53 mg/kg.

### Experimental procedures

After the period of acclimatization to the laboratory environment, the animals were divided into the following experimental groups: Control (CTRL), DMSO, OTA (1.38, 2.77, and 5.53 mg/kg). Allocation to experimental groups followed randomization procedures with a computerized random number generator (random.org) and the procedure was performed by researchers blinded to the experimental group. The drugs for each experimental group were administered at the beginning of the experiment (at 0 hours) by intraperitoneal injections and the control group received saline. Briefly, the intraperitoneal injections were performed using a Hamilton Microliter^™^ Syringe (701 N 10 μL SYR 26 s/2” /2) x Epidurakatheter 0.45 × 0.85 mm (Perifix^®^-Katheter, Braun, Germany) x Gingival Needle 30G/0.3 × 21 mm (GN injecta, SP, Brazil). The injection volume was 1 μL/100 mg of animal weight. The animals were previously anesthetized by immersion in a solution of tricaine (300 mg/L) until loss of motor coordination and reduced respiratory rate. After the anesthesia, the animals were placed in a sponge soaked in water exposing the abdomen and the needle was gently inserted parallel to the spine in the abdomen’s midline posterior to the pectoral fins. This procedure was conducted in approximately 10 seconds (Fig. 1A) (Bertelli et al., 2021).

**Figure 1.**
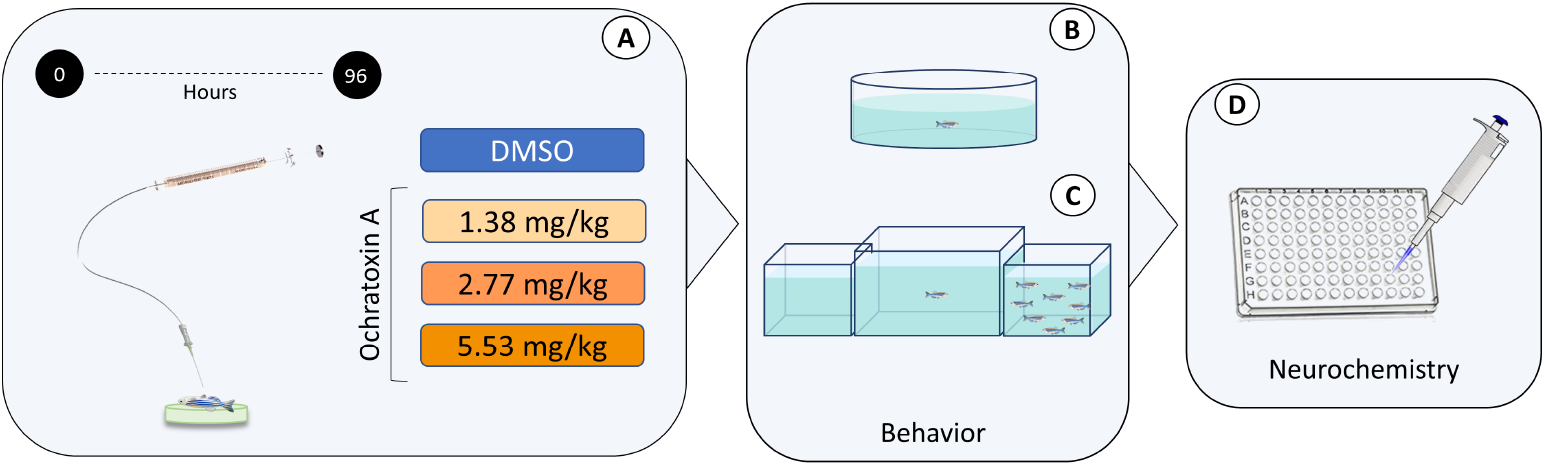
Experimental design.

Following drug administration, the fish returned to their respective experimental tanks and remained for 96 hours. The housing and feeding conditions of the experimental tanks were the same described for the laboratory housing tanks. After 96 hours of exposure, the animals were individually submitted to the open tank test (OTT). After this, the animals returned to the experimental tank and remained for 24 hours. Then, the animals were submitted to the social interaction test (SIT). Immediately after the SIT, the animals were euthanized, and the brains were dissected and homogenized for the neurochemical assays of the parameters associated with oxidative status. The neurochemical parameters analyses were: thiobarbituric acid reactive substance (TBARS), non-protein thiol (NPSH), glutathione peroxidase (GPx), glutathione-S-transferase (GST), and glutathione reductase (GR). The sex of the animals was confirmed after euthanasia by dissecting and analyzing the gonads.

### Open tank test (OTT)

The OTT consists of a white circular arena (24 cm diameter, 8 cm height, and 2 cm water level). In this test, the animals were placed in the center of the arena and the behavior was individually recorded for 10 min (Fig. 1B). The videos were obtained from an upper view and for the analyses, the arena was virtually divided into two zones: center and periphery (Benvenutti et al., 2020). The following parameters were quantified using ANY-Maze software (Stoelting Co., USA): distance, crossings, absolute turn angle, mean speed, freezing episodes, and time-freezing duration.

### Social interaction test (SIT)

In the SIT, fish were placed individually in a central tank (30 x 10 x 15 cm) flanked by two identical tanks (15 x 10 x 13 cm) and filmed from a frontal view for 7 min (Fig. 1C). One of the two tanks positioned beside the central tank (test tank) contained only water (neutral stimulus), and the other contained 10 zebrafish (social stimulus). All tanks were filled with water at a level of 10 cm and in the same conditions. The side of the social stimulus tank was counterbalanced to avoid any eventual bias (Benvenutti et al., 2020). The analyzes were carried out with the aid of the ANY-Maze software (Stoelting Co., USA), and for that, the test tank was virtually divided into three equal vertical zones (interaction, middle and neutral). The interaction zone was considered to be next to the tank that contained the social stimulus, while the neutral zone was considered to be next to the neutral stimulus. Animals were placed in the middle zone and had 2 min to habituate to the tank test. After this, the behavior was analyzed for 5 min. The parameters quantified were distance, number of crossings, and interaction time.

### Neurochemical analysis

Following the behavioral tests, the animals were euthanized by hypothermic shock (2°-4 °C) and decapitation. The brains samples were then collected to evaluate the oxidative status (Fig. 1D). For each sample, a pool of 4 brains was used (n = 6) and mixed with 600 μL of phosphate-buffered saline (PBS, pH 7.4, Sigma-Aldrich). The homogenate was centrifuged at 2400 g for 10 min at 4 °C and the supernatants were collected for the analyses of the following parameters: lipid peroxidation (TBARS) (Sachett, 2020), non-protein thiols (NPSH) (Sachett, 2020), and glutathione-S-transferase (GPx) (Sachett, 2021a), glutathione reductase (GR) (Sachett, 2021b) and glutathione-S-transferase (GST) (Habig and Jakoby, 1981).

### Statistical analysis

The sample size was calculated with power analysis using G*Power 3.1.9.2 for Windows. Normality and homogeneity of variances were confirmed for all data sets using D’Agostino-Pearson and Levene tests, respectively. The student’s t-test was performed to compare control and DMSO groups. One-way ANOVA followed by Tukey’s post hoc test was used for the analyses. For behavioral data, the outliers were identified based on distance traveled using the ROUT statistical test (GraphPad^®^ software) and were removed from the analyses. This resulted in 3 outliers (2 animals from the DMSO group and 1 animal from OTA 2.77 mg/kg group) removed from the OTT and 3 outliers (1 animal from the DMSO group, 1 from the 2.77 mg/kg group, and 1 from the 5.53 mg/kg group) removed from the SIT. The tank and sex effects were tested in all comparisons; The data were expressed as mean ± standard deviation (S.D.). Differences were considered significant at p<0.05.

## RESULTS

DMSO did not show important modulation on behavior (Supplementary material 1) or induce oxidative damage (Supplementary material 2) compared with control sodium chloride. Therefore, we have used DMSO as a control group.

### Open tank test

Fig. 2 shows the acute effects of OTA in adult zebrafish in the open tank test. There was a significant decrease in the distance (Fig. 2A, p = 0.0105), absolute turn angle (Fig. 2C, p = 0.0090), mean speed (Fig. 2D, p = 0.0110) and an increase in time-freezing (Fig. 2F, p = 0.0052) at the 1.38 mg/kg dose, indicating locomotor damage. The parameters of crossings and freezing episodes did not show significant changes in any dose.

**Figure 2.**
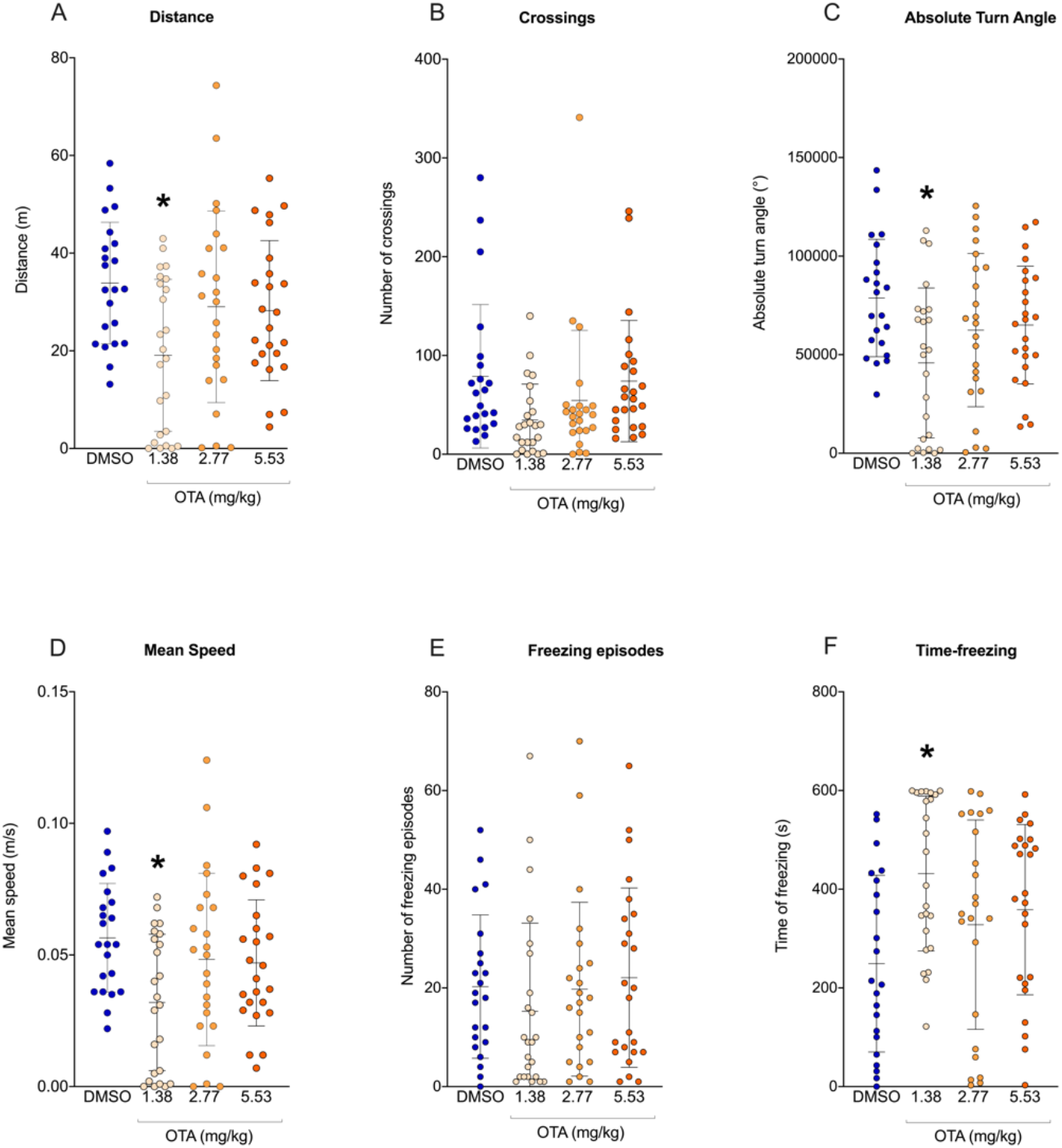
Effects of OTA in the open tank test. (A) distance, (B) crossings, (C) absolute turn angle, (D) mean speed, (E) freezing episodes, and (F) time-freezing. Data are expressed as mean ± standard deviation (S.D.). n= 21-24. One-way ANOVA followed by Tukey’s post hoc test. *p < 0.05.

### Social interaction test

Fig. 3 shows the acute effects of OTA on adult zebrafish at the SIT. OTA, in the tested doses, did not alter social behavior in any of the analyzed parameters.

**Figure 3.**
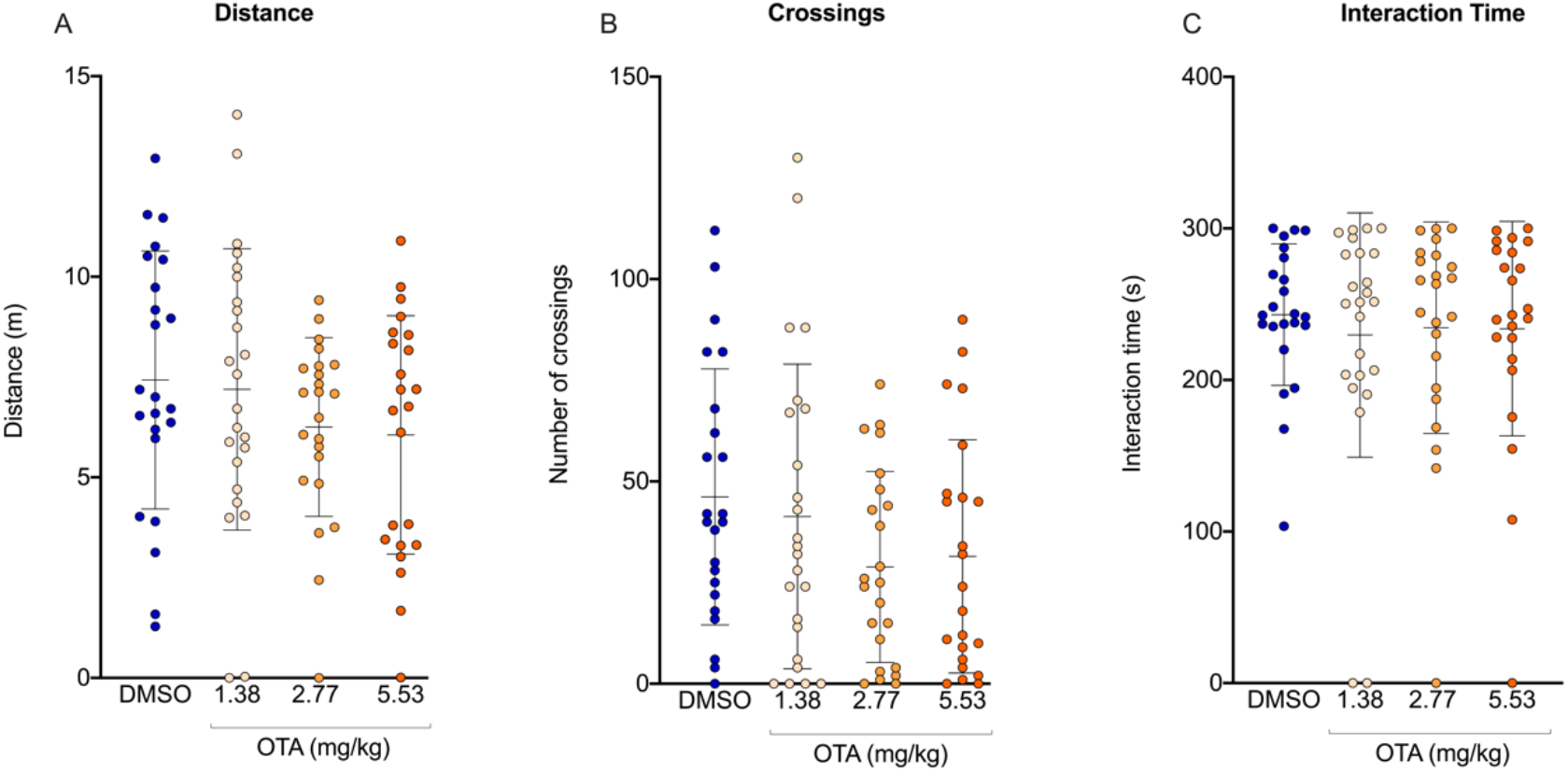
Effects of OTA in the social interaction test. (A) distance, (B) crossings, (C) interaction time. Data are expressed as mean ± standard deviation (S.D.). n= 21-24. One-way ANOVA.

### Neurochemical analysis

Fig. 4 shows the effects of OTA on neurochemical parameters. OTA at 1.38 mg/kg increased the GPx (Fig. 4C, p < 0.0001), GST (Fig. 4D, p < 0.0001), and GR (Fig. 4E, p = 0.0397) activities. The intermediate dose of 2.77 mg/kg decreased NPSH levels (Fig. 4B, p = 0.0006) and increased GPx (Fig. 4C, p = 0.0016) and GST (Fig. 4D, p = 0.0146) activities. The dose of 5.53 mg/kg increased GPx (Fig. 4C, p < 0.0001) and GR (Fig. 4E, p = 0.0238) activities.

**Figure 4.**
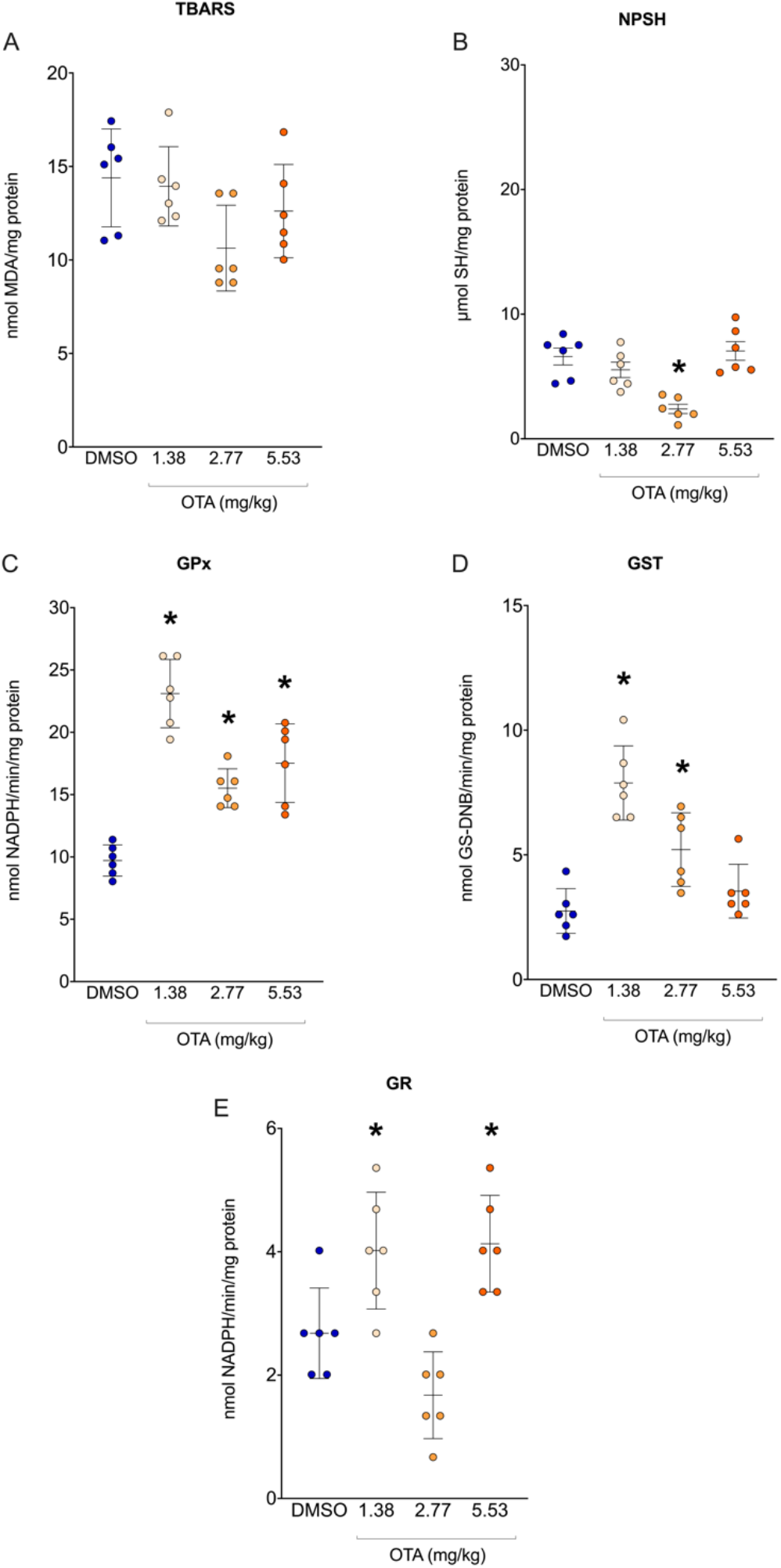
Effects of OTA in neurochemical parameters. (A) TBARS, (B) NPSH, (C) GPx, (D) GST, and (E) GR. Data are expressed as mean ± standard deviation (S.D.). n= 6. One-way ANOVA followed by Tukey’s post hoc test. *p < 0.05.

## DISCUSSION

This study showed the deleterious effects of ochratoxin A in adult zebrafish. Briefly, the toxin decreased the total distance traveled, average speed, absolute turn angle and increased the freezing time. However, in the social interaction test, there were no behavioral changes in the evaluated parameters. Neurochemical analysis showed that the compound was able to generate alterations in the oxidative status that were not enough to alter TBARS but were able to trigger the oxidative defense system.

There are little data in the literature on the behavioral effects of OTA exposure not only in fish but this lack of information also extends to other animals. In zebrafish larvae, OTA decreased the animals’ swimming speed but did not change parameters of distance and time spent active (Khezri et al., 2018). In rodents, it was shown that OTA injected intraperitoneally was able to cause behavioral changes in gait analysis, spontaneous activity, cylinder test, and pole test, similar to Parkinsonian symptoms that were stabilized with the use of L-Dopa (Bhat et al., 2018).

In our study, the interference of OTA on locomotion parameters in zebrafish was shown in the open tank test similar to the results previously cited in other models. A possibility for these findings could be the link between locomotion and the nigrostriatal pathway that has already been reported to be affected by OTA in rodent models (V. Sava et al., 2006; V Sava et al., 2006b). However, there were no changes in social interaction parameters. The social behavior in zebrafish presents a schooling cohesion that aims to search for food, escape from predators and reproduce (Pitcher, 1993). Thus, being a model closely linked to social functions, the zebrafish has been extensively studied for this type of behavior (Buske and Gerlai, 2011; Dreosti et al., 2015; Scerbina et al., 2012). However, precisely because socialization is genetically preserved and has an ontogenic nature in zebrafish, it may be a parameter less vulnerable to milder modulations such as those shown in this study, since the lowest concentration used was about 25% of the LD50 established in another model. Another aspect to be considered is related to the cues provided by the apparatus since previous studies have already demonstrated the multifactorial character of social behavior in zebrafish being linked to visual cues (Engeszer et al., 2007), olfactory cues (Gerlach et al., 2007) and also sensitive to alarm substances released by co-specifics (Canzian et al., 2017). The apparatus used in this study, however, only allowed the visual cues to be transmitted to the animal, so it is uncertain to say what the effects of OTA would be under other parameters involved in the animal’s social behavior.

With increasing global concern about the spread of mycotoxins, the effect of these compounds on oxidative stress parameters has become a very debated issue (da Silva et al., 2018; Mavrommatis et al., 2021), with emphasis on ochratoxin A (Sorrenti et al., 2013; Tao et al., 2018). OTA can interact with peroxidases that produce a phenoxyl radical from OTA. Glutathione (GSH) is capable of turn the phenoxyl radical into OTA again by forming a superoxide anion radical (O_2_^•–^) that results in hydrogen peroxide (H_2_O_2_). H_2_O_2_ by Fenton reaction produces a hydroxyl radical (OH^•^) that is responsible for oxidative damage (Adlouni et al., 2000). Another common pathway for OTA is the formation of an OTA–Fe^3+^ complex that is reduced in OTA-Fe^2+^ by cytochrome P450 resulting in OH^•^ (Rahimtula et al., 1988). Several studies report the imbalance of oxidative status caused by the compound. In zebrafish larvae, there was the formation of reactive oxygen species (ROS) proportional to the increase in OTA concentration (Tschirren et al., 2018). A study with tambaqui (*Colossoma macropomum*), a freshwater fish, found an increase in the ROS and lipid peroxidation in the animal’s muscles, as well as a decrease in the activity of antioxidant enzymes superoxide dismutase (SOD) and GPx (Baldissera et al., 2020). Similarly, an increase in lipid peroxidation and antioxidant enzymes activity catalase (CAT) and GR was seen with a decrease in SOD activity and GSH levels in the brain, kidney, and liver of rats (Nogaim et al., 2020). A study found an increase in ROS formation, lipid peroxidation, and decreased GSH levels in kidney cells (Lee et al., 2018). However, studies with birds have shown that in long-term exposure antioxidant defenses can increase against oxidative imbalance, especially the glutathione redox system (Fernye et al., 2021; Kövesi et al., 2019). Also, a study with *Caenorhabditis elegans* shown an increase in the expression of SOD and CAT in wine containing OTA (Schmidt et al., 2020). These studies collaborate with our results which showed that, in adult zebrafish, there was an increase in enzyme defenses with an elevation of GPx, GR, and GST, especially at the lowest dose. In the intermediate dose, there was no increase in GR as occurred in the other doses, which is consistent with the decrease in GSH levels (NPSH) in this group since GR is responsible for the recycling of glutathione, which is essential for the maintenance of antioxidant levels. The increase in GPx under these conditions indicates an attempt to control a possible increase in reactive oxygen species since GPx reduces H_2_O_2_ through the GSH oxidation, something quite common to occur in OTA exposures as mentioned in previous studies. The increase of GST activity also indicated an increase in OTA metabolization and elimination since GST catalyzes the conjugation of the reduced form of glutathione to xenobiotic substrates for detoxification. Likewise, this activation of defenses prevented the increase of ROS levels and consequently avoiding lipid peroxidation (TBARS levels) (Dasuri et al., 2013; Gandhi and Abramov, 2012).

Despite the zebrafish being a model used for decades in research in several areas, many gaps still exist about the model, especially in the area of toxicology. In recent years there has been a considerable increase in studies in this field due to initiatives to standardize this type of analysis in fish (Gonçalves et al., 2020), including the OECD protocols (OECD Guidelines for the Testing of Chemicals, 1992). However, for adult animals, the methodologies tend to be limited to direct exposure to the animals’ water, which is not suitable for all protocols. In the case of OTA, the formulation of the compound and the difficulty in storing or disposing of waste made this type of exposure impracticable so the intraperitoneal injection standardized in the laboratory was chosen. The use of intraperitoneal injection to assess the effects of OTA has already been used in other models, being effective in detecting deleterious effects on the metabolism mechanism in rats (Størmer et al., 1985) and neurotoxicity in the development of mice (Miki et al., 1994; Tamaru et al., 1988). In fish, OTA was injected peritoneally into rainbow trout (*Salmo gairdneri*) acutely (96 hours) for toxicological evaluation by histology and determination of LD50 (5.53 mg/kg) (Doster et al., 1974). However, these data were never detailed in other species and the use of zebrafish to evaluate the effects of OTA remained limited with little information regarding the effects of the toxin in this species.

Due to these important gaps in the literature, another point to be clarified is the dose-response reaction of zebrafish against OTA. In this study, the doses that were more behaviorally and neurochemically reactive were the lowest doses, with the highest dose changing few parameters in oxidative status. Thus, in this study, we speculate that OTA showed a hormetic effect in adult zebrafish. Hormesis is a biphasic dose-response characterized by stimulation at low doses and inhibition at high doses (Calabrese and Baldwin, 2002). For OTA, this type of curve has already been reported in an *in vitro* study (Li et al., 2014), however, this is the first time that this behavior has been seen in an *in vivo* model. A biphasic curve can indicate the biological plasticity of the target organism (Calabrese and Mattson, 2011), and the zebrafish is a widely studied model precisely because of its capacity for neuroplasticity and regeneration (Cosacak et al., 2015; Ghosh and Hui, 2016). Thus, it is possible that the hormetic behavior of OTA, in this case, is linked to the animal’s biological characteristics. Moreover, hormetic curves often occur with endocrine disruptors (Vandenberg et al., 2012) and other studies have demonstrated the potential of OTA to interfere with hormone production (Frizzell et al., 2013; Woo et al., 2013). For all these reasons, toxicological results for low doses should not be ignored.

## CONCLUSION

Although concern about controlling OTA levels is increasing, more efforts are still needed. For this, understanding the effects of the toxin on organisms is essential. This study demonstrated the potential that the toxin has for causing deleterious effects in adult zebrafish through behavioral change and neurochemical modulation, however, more studies are needed to elucidate the compound’s mechanism of action and its effects on other organisms to further contribute to the field of toxicology and environment.

## ACKNOWLEDGMENTS

We thank the Coordenação de Aperfeiçoamento de Pessoal de Nível Superior - Brasil (CAPES), Conselho Nacional de Desenvolvimento Científico e Tecnológico (CNPq, proc. 303343/2020-6), and Pró-Reitoria de Pesquisa (PROPESQ) at Universidade Federal do Rio Grande do Sul (UFRGS) for funding.

## AUTHOR CONTRIBUTIONS

All authors had full access to all the data in the study and take responsibility for the integrity of the data and the accuracy of the data analysis. Conceptualization, J.V. and A.P.; Methodology, J.V., A.S., M.M., L.M.B., A.P.; Investigation, J.V., A.S., M.M., L.M.B.; Formal Analysis, J.V., A.S., M.M., L.M.B., A.P.; Resources, A.P.; Writing - Original Draft, J.V.; Writing - Review & Editing, J.V., A.S., M.M., L.M.B., A.P.; Supervision, A.P.; Funding Acquisition, A.P.

## COMPETING INTERESTS

The authors declare no competing interests.

## SUPPLEMENTAL MATERIAL

**Supplementary material 1.**
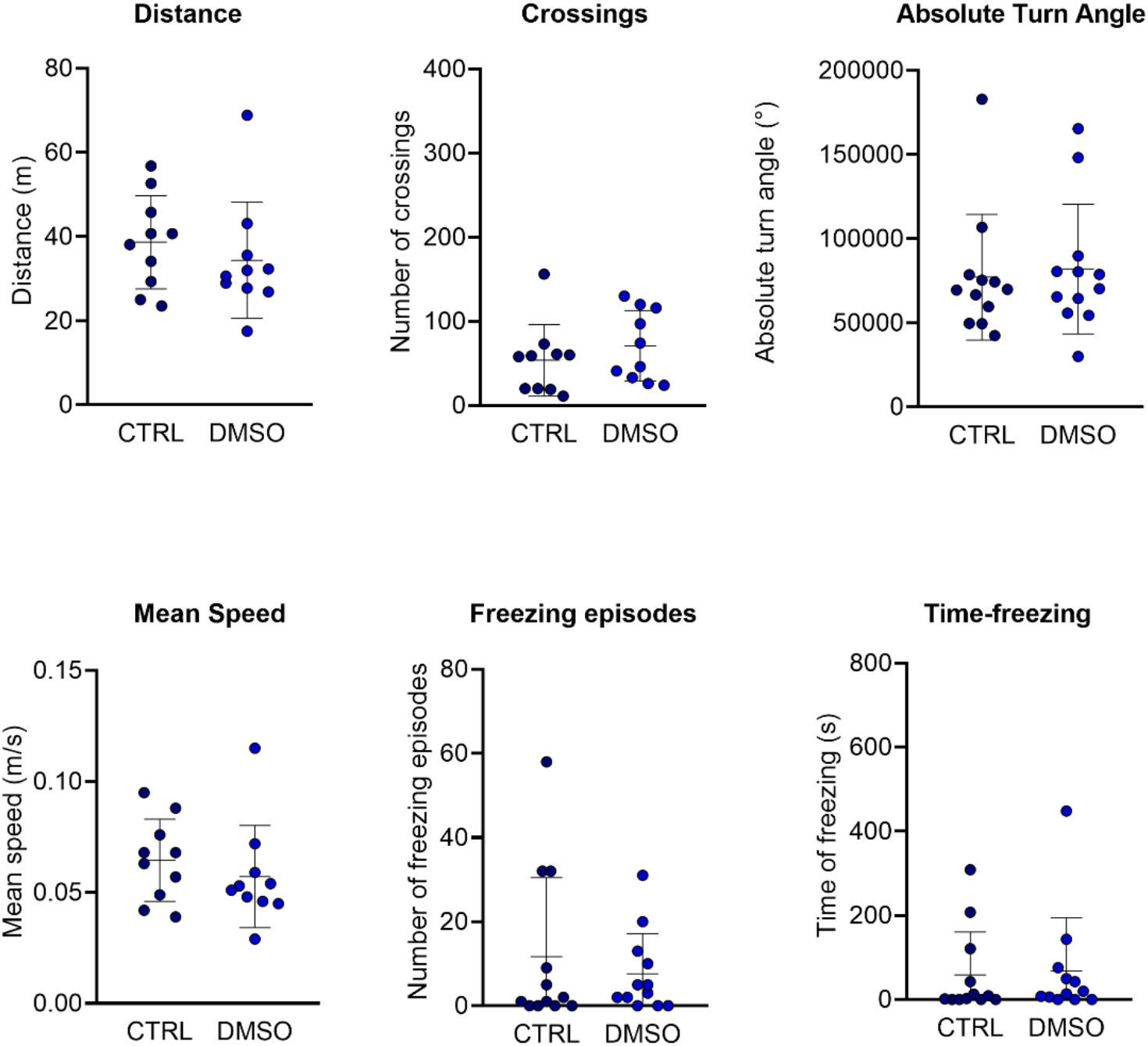
Comparison between CRTL (sodium chloride) vs. DMSO (dimethyl sulfoxide) on behavior parameters at open tank test. Data are expressed as mean ± standard deviation (S.D.). n= 10. Student’s t-test. *p < 0.05.

**Supplementary material 2.**
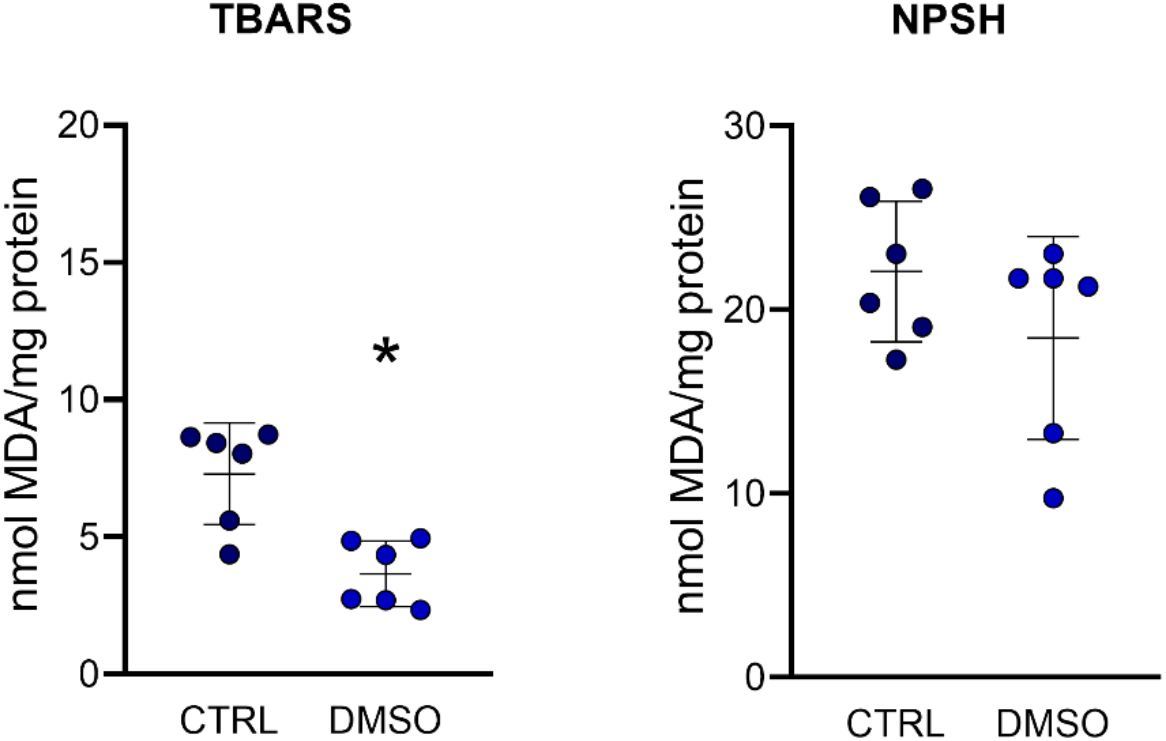
Comparison between CRTL (sodium chloride) vs. DMSO (dimethyl sulfoxide) on neurochemical parameters. Data are expressed as mean ± standard deviation (S.D.). n= 6. Student’s t-test. *p < 0.05.

## Notes

### Competing Interest Statement

The authors have declared no competing interest.

